# Human type-I interferon omega holds potent antiviral properties and promotes cytolytic CD8^+^ T cell responses

**DOI:** 10.1101/2025.02.13.638030

**Authors:** Hoang Oanh Nguyen, Patricia Recordon-Pinson, Marie-Line Andreola, Laura Papagno, Victor Appay

## Abstract

The type-I interferon family is well known for its critical role in innate immunity. It comprises several members, among which IFN-α2 and IFN-β are the most extensively studied, with important antiviral and immune-modulatory functions. Recent findings linking autoantibodies against type-I interferons to severe COVID-19 suggest a potential role for IFN-ω in combating SARS-CoV-2 infection. However, little is known about human IFN-ω, as most research on this interferon has been conducted in feline models. Here, we demonstrate that human IFN-ω is secreted at levels comparable to those of IFN-α2 or IFN-β upon stimulation with inflammatory agonists and triggers a robust antiviral response, inhibiting SARS-CoV-2 infection *in vitro*. Moreover, IFN-ω enhances the effector functions of antigen-specific CD8^+^ T cells primed *de novo* from healthy donor cells, highlighting its capacity to promote strong cellular immunity. Our results position IFN-ω as a key member of the type-I interferon family, with promising potential for therapeutic and vaccine applications.

**Author Summary:** Type-I interferons are pleiotropic cytokines, including IFN-α and IFN-β, which are well known for their antiviral activities. Here, we report on the functional characteristics of human IFN-ω, a neglected member of the type-I IFN family, which has primarily been studied in feline models. We show here that human IFN-ω induces intense downstream signaling in cells resulting in the upregulation of antiviral genes, and efficient restriction of SARS-CoV-2 replication. IFN-ω also promotes the acquisition of strong cytolytic functions by antigen primed CD8^+^ T cells. Overall, our findings portray human IFN-ω as a major antiviral molecule, similar to the well-studied and highly effective IFN-α_2_. The secretion of IFN-ω upon infection is likely to be crucial for effective control of multiple viruses, advocating for its use in therapeutic approaches in humans.

## Introduction

Type-I interferons (IFNs-I) are known for their striking antiviral capacity^1^ and role in regulating both innate and adaptive immune responses^2^. These cytokines can interfere at many levels of the viral replication cycle within cells, thereby preventing viral infection and propagation^3,4^. Hence, IFNs-I have been the center of interest in many therapeutic applications and preventive medicine^5,6^. The human IFNs-I family comprises up to 16 members, including IFN-α and its many subtypes, IFN-β, IFN-ε, IFN-κ and IFN-ω. All share a relatively homologous structure and bind to the ubiquitously expressed transmembrane receptor composed of two chains, IFNAR1 and IFNAR2^7^, although with different affinities. IFN-α_2_ and IFN-β have strong affinity for IFNAR, in contrast to IFN-ε, which has a reduced affinity for the IFNAR2 receptor chain and thus a weaker activity^8^. Upon binding to their cognate receptors, IFNs-I activate the canonical JAK/STAT1/IRF9 cascade, which in turn upregulate a large network of Interferon-stimulated genes (ISGs). These genes orchestrate distinct biological functions of IFNs-I that were thoroughly described based on the characteristics of the prototypical IFN-α_2_ and IFN-β^3,4^.

To date, little research has been conducted on IFN-ω. This interferon, discovered in 1985 by Hauptmann et al^9^, has been identified in humans and several animal species, but not in mice. IFN-ω was thus mainly characterized in feline, where it displays strong anti-viral efficacy^10-13^. Hence, recombinant feline IFN-ω has been registered and widely prescribed for the treatment of viral infections in cats in many European countries^14^. However, its functional properties and potential therapeutic benefits have been poorly investigated in the human setting. Recombinant IFN-ω was shown to be well tolerated in HCV infected patients and efficiently reduce HCV RNA levels when combined with ribavirin^15-17^. A recent study suggests a role of IFN-ω against SARS-CoV-2 infection in children and young adults^18^. Moreover, high levels of auto-antibodies against IFN-ω have been reported to be associated with high mortality rate in severe COVID-19 patients^19,20^. These studies support a potential role of IFN-ω against virus infections in humans and call for an in-depth characterization of its antiviral properties in a human setting. Here we compared the properties of IFN-ω with those of IFN-α_2_ across several assays to assess its antiviral and immunomodulatory potency using human cell lines and PBMCs.

## Results

### Human monocytes and plasmacytoid DCs produce high levels of IFN-ω

We first aimed to assess the bioavailability of IFN-ω. To this end, we stimulated human PBMCs with different proinflammatory agents that mimic virus-induced activation, and measured the secretion of IFN-ω in comparison with IFN-α_2_ and IFN-β after 24 hours. We used cGAMP and CpG-A, the ligands for STING and TLR9 respectively, known to induce strong IFN-I responses. For comparison, we also used the TLR4 and TLR8 agonists, LPS and ssRNA40, which induce strong inflammatory but weak IFN responses. Both cGAMP and CpG-A stimulation yielded high levels of IFN-ω, in comparable range to IFN-α_2_ and IFN-β. (**Figure 1A**). On the other hand, ssRNA40 was capable of inducing the production of IFN-β and IFN-ω, but not that of IFN-α_2_. As expected, LPS elicited little to no secretion of all types of IFN-Is.

**Figure 1.**
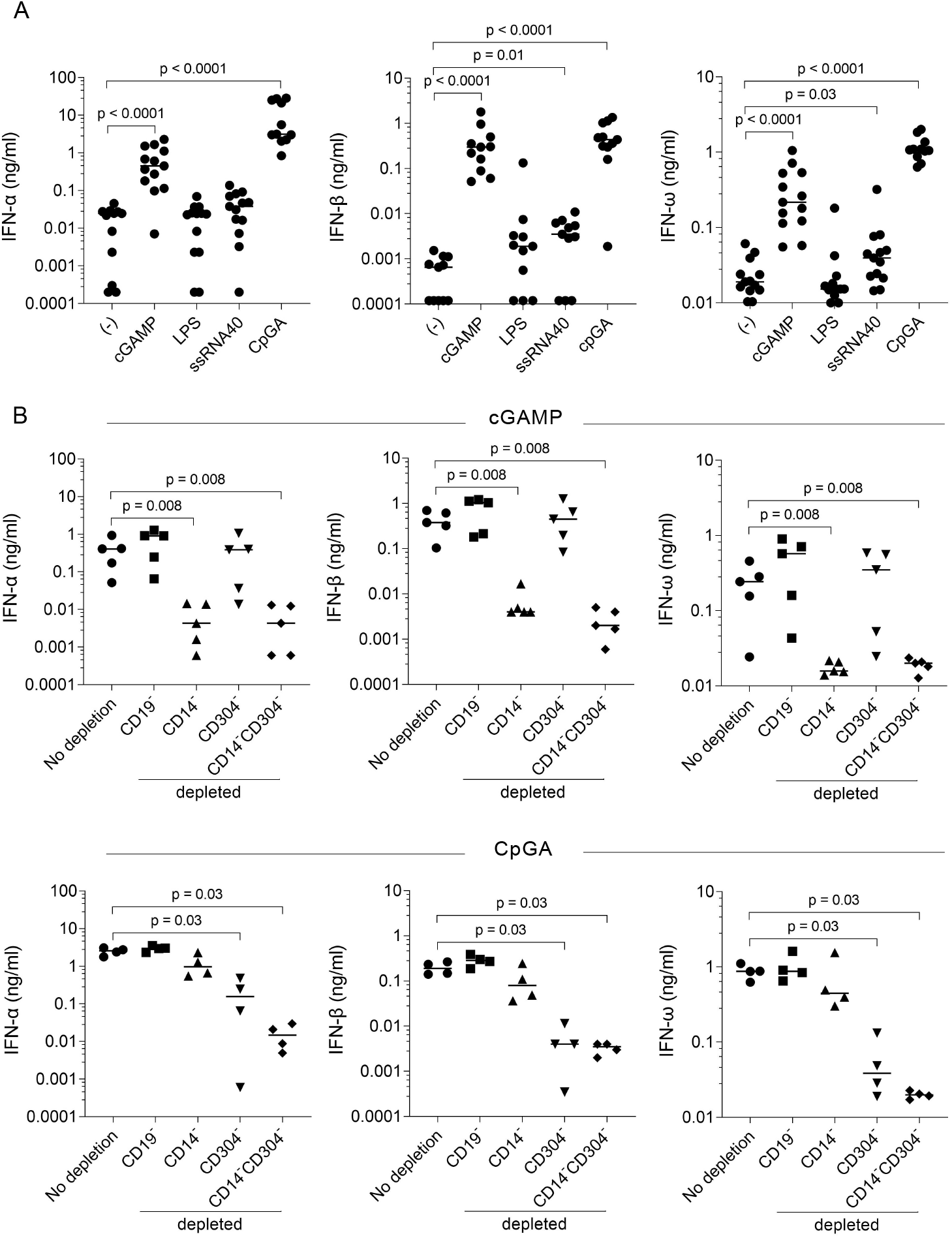
Monocytes and plasmacytoid dendritic cells are the principal source of IFN-ω. **A**. Secretion of IFN-α, IFN-β and IFN-ω by human PBMCs (n=13) upon stimulation with either cGAMP (10 μg/ml), LPS (100 ng/ml), ssRNA40 (500 ng/ml) or CpG-A (2 μg/ml) for 24 hours. Levels of type-I IFNs (α, β, ω) were quantified by ELISA in cell-free supernatants. Each dot represents one donor. **B**. Levels of IFN-α, IFN-β and IFN-ω in the supernatants of human PBMCs (n=5) depleted for CD19^+^ B cells, CD14^+^ monocytes, or CD304^+^ pDCs, upon stimulation with either cGAMP (10 μg/ml) or CpG-A (2 μg/ml) for 24 hours. Each dot represents one donor. Horizontal bars indicate median value. Differences between groups were tested using the non-parametric Mann Whitney test.

To investigate which cell types were responsible for IFN-ω production, we selectively depleted CD14^+^ monocytes, CD304^+^ plasmacytoid dendritic cells (pDCs) or CD19^+^ B cells from total PBMCs and subsequently stimulated the remaining cells with cGAMP or CpG-A. pDCs and monocytes are known to be the main producers of IFN-α upon stimulation with CpG-A and cGAMP respectively^21^. Depletion of monocytes resulted in the loss of IFN-ω secretion in response to cGAMP, whereas depletion of pDCs resulted in the loss of IFN-ω secretion in response to CpG-A (**Figure 1B**). Accordingly, simultaneous depletion of these two populations completely abrogated IFN-ω production with either CpG-A or cGAMP. Similar observations were made for IFN-α and IFN-β production. In contrast, the depletion of B cells did not affect IFNs-I levels. Monocytes and pDCs are therefore the main source of IFN-ω production, which is determined by the type of ligands used for stimulation. The profile and levels of IFN-ω production are similar to those of IFN-α and IFN-β making IFN-ω equivalent to its two sister molecules in terms of bioavailability.

### IFN-ω induces the STAT1/IRF9 pathway and has potent antiviral activity

We next investigated downstream signaling and antiviral activity of IFN-ω. A dual reporter Jurkat cell line was used to study signaling through NF-kB and IRF pathways simultaneously in the presence of different IFNs-I or Poly I:C, a TLR3L ligand, used as positive control. All tested IFNs-I, except the low affinity binder IFN-ε, induced expression of SEAP reporter protein, indicating activation of the STAT1/IRF9 pathway (**Figure 2A**). In contrast, NF-kB signaling remained inactive as low levels of Lucia reporter protein were observed in the presence of IFNs-I. In dose response assays, IFN-ω performed as efficiently as IFN-α_2_ or IFN-β in inducing the IRF pathway (**Figure 2B**). These results indicate that IFN-ω binds to IFNARs with high affinity, thus activating efficiently the classic IFN/IRF pathway. Since antiviral activity is a distinctive characteristic of IFNs-I, we explored whether IFN-ω displays similar antiviral potencies as to IFN-α_2_. To this end, we monitored the kinetic expression of mRNAs encoding for proteins associated with the cellular antiviral response, including MX1, OSA2, ISG15, IFI27, IFI2 and MX2, in Calu-3 cells, a pulmonary human cell line, upon exposure to IFN-ω^22^. Significant upregulation of these genes was observed in the presence of IFN-ω, similarly to IFN-α_2,_ while IFN-ε triggered no antiviral gene profile (**Figure 2C**).

**Figure 2.**
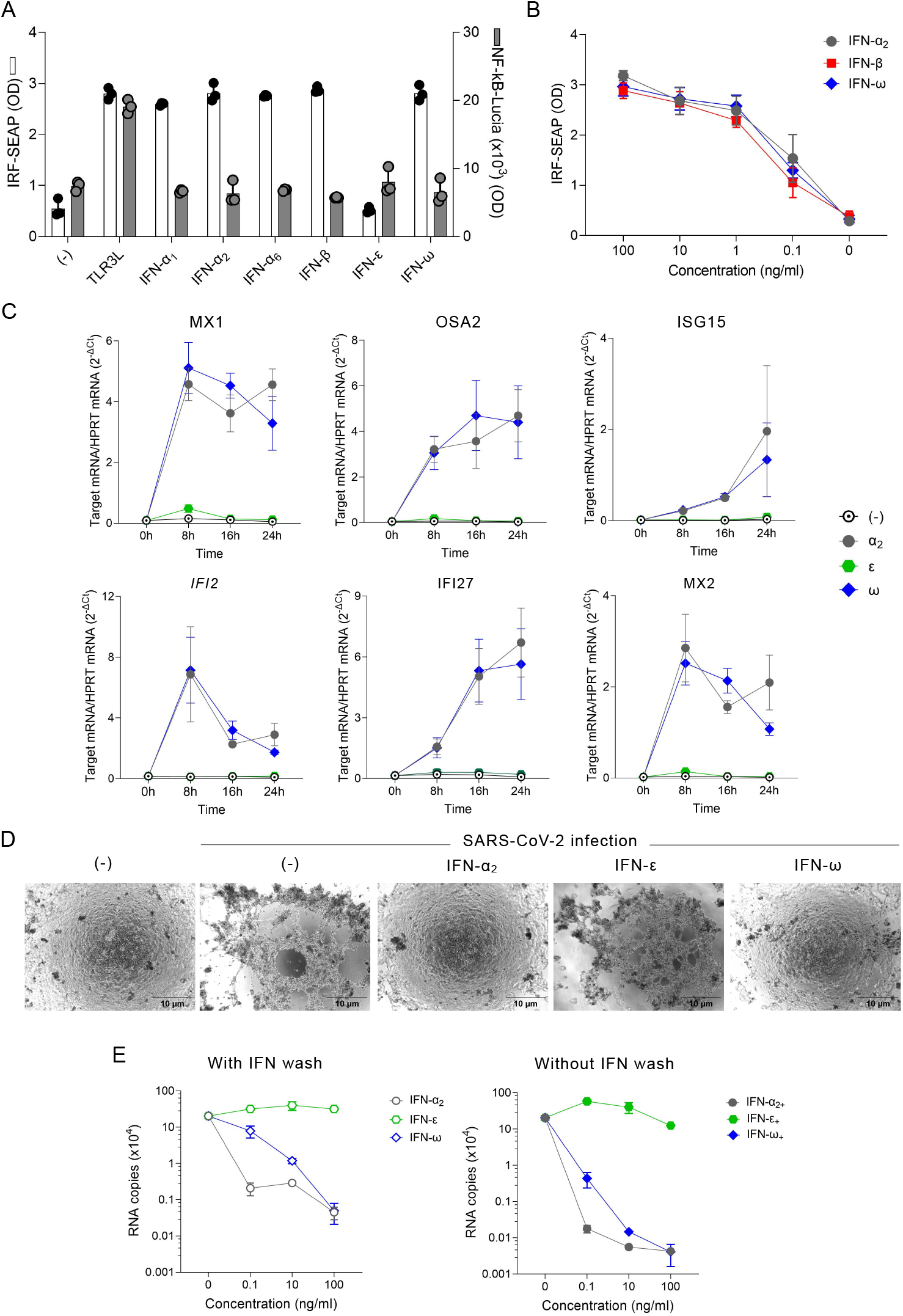
IFN-ω triggers a strong antiviral program in human cells. **A**. Induction of IRF and NF-κB signaling pathways in human cells by type-I IFNs. The Jurkat-Dual™ cell line was stimulated with the TLR3 ligand Poly(I:C) at 5 µg/ml or different IFNs-I (α_1_, α_2,_ α_6,_ β, ε or ω) at 100ng/ml for 24 hours. IRF or NF-κB activation was evaluated by measuring the levels of SEAP and Lucia luciferase in cell-free supernatants using QUANTI-Blue™ Solution or Luc™ respectively. Data are expressed as the mean ± SEM of the optical density values of SEAP (left y axis) and Lucia levels (right y axis) from three independent experiments. **B**. IRF activation was evaluated by measuring the levels of SEAP in supernatants of Jurkat-Dual™ cells after 24h stimulation with IFN-α_2_, β or ω at indicated concentrations. **C**. Expression of mRNAs encoding proteins associated with the cellular antiviral response in Calu-3 cells at different times upon exposure to IFN-α_2_, ε or ω (100 ng/ml). RNA expression levels were evaluated by Real-Time PCR. Data are expressed as mean ± SEM (n = 3) of 2^−ΔCt^ relative to housekeeping mRNA (HPRT). **D**. Cellular morphology of uninfected or SARS-CoV-2 infected Calu-3 cells (MOI = 0.1) in the absence or in the presence of type-I IFNs (100 ng/ml) using an electron microscope. **E**. Quantification of SARS-CoV-2 replication in Calu-3 cells pre-treated with the indicated doses of IFN-α_2_, ε or ω for 16 hours and infected with SARS-CoV-2 (MOI = 0.1) for 2 hours. Viral suspension was then replaced with medium alone (left panel) or medium containing the respective IFNs (right panel). Viral RNA copies were quantified using RT-PCR. Data are expressed as the mean ± SEM of viral RNA copies from two independent experiments.

We next set up a viral infection model by infecting Calu-3 cells with SARS-CoV-2. Calu-3 cells were treated with different concentrations (0.1, 10, 100 ng/ml) of IFN-ω, IFN-α_2_ or IFN-ε for 16 hours before viral infection to ensure that both early and late ISGs would be fully induced^22^. Where indicated, the cells were maintained with IFNs-I after the addition of viral suspension. The antiviral effect of IFN-ω was assessed based on Calu-3 morphology and viral replication after infection. Pretreatment with IFN-ω efficiently protected Calu-3 cells against SARS-CoV-2 invasion, characterized by a preserved cell morphology (**Figure 2D**) and low levels of viral RNA copies (**Figure 2E**). The continual presence of IFN-ω at high dose (100 ng/ml) before and after the infection fully inhibited SARS-CoV-2 replication. Of note, IFN-α_2_ outperformed IFN-ω in preventing viral replication, at low doses (i.e. 0.1 ng/ml). In contrast, IFN-ε presented minimal to no suppressive effects on SARS-CoV-2 infected Calu-3 cells, consistent with its previously reported weak antiviral characteristic^23-25^. The findings indicate that the interaction of IFN-ω with IFNARs induces intense downstream signaling along with high antiviral profile defined by the upregulation of MX1 and OSA2 mRNA levels, similarly as to IFN-α_2_, resulting in efficient restriction of SARS-CoV-2 replication and protection of Calu-3 cells from cytopathic effects. IFN-ω possesses therefore potent antiviral properties and its secretion upon infection is likely crucial for effective viral clearance.

### IFN-ω bridges innate and adaptive immunity by potentiating the effector functions of antigen-primed CD8^+^ T cells

IFN-α_2_ and IFN-β are essential for the differentiation of naive CD8^+^ T cells into effector cells by tuning a complex network of STAT molecules, including STAT4, whose expression is positively associated with optimal cell expansion and function, and STAT1, which is associated with antiviral genes and cell inhibition^26,27,28^. We thus compared the phosphorylated forms of STAT4 and STAT1 in subsets of CD8^+^ T cells treated with IFN-ω, IFN-α_2_ or IFN-β. For these experiments, a concentration of 100 ng/ml of IFNs-I was used, as this dose yielded the strongest signaling response. Upon stimulation with IFNs-I, STAT1 and STAT4 were rapidly induced in naive and memory CD8^+^ T cells and the activation patterns of both pSTATs overlapped between the three IFNs-I over time **(Figure 3A)**. These data demonstrate that IFN-ω resembles IFN-α_2_ in terms of STAT regulation and that both STAT1 and STAT4 phosphorylation occur concurrently following IFNs-I exposure.

**Figure 3.**
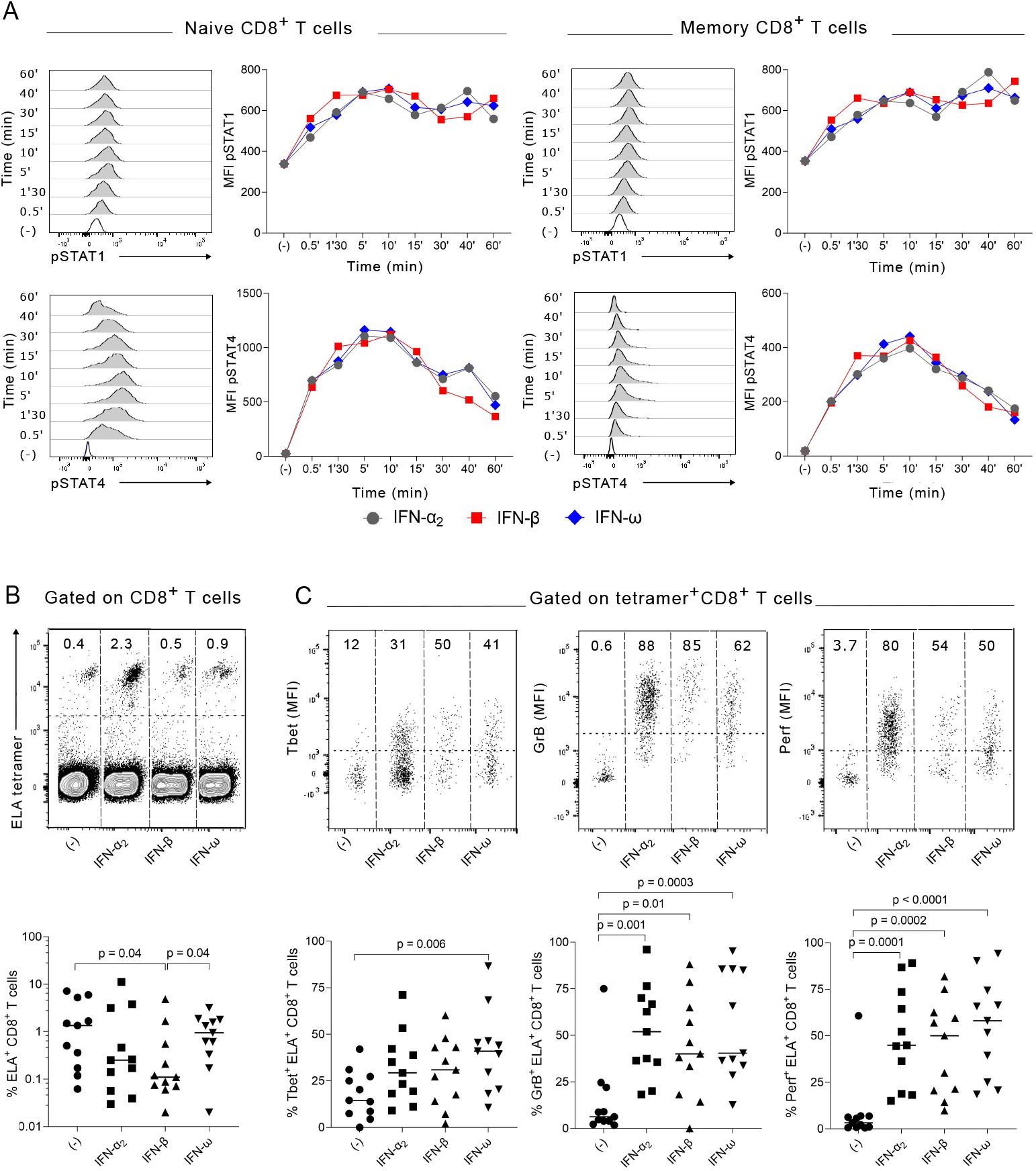
IFN-ω triggers STAT1 and STAT4 activation and enhances effector functions of antigen primed CD8^+^ T cells. **A**. STAT1 and STAT4 phosphorylation in CD8^+^ T cell subsets stimulated with type-I IFNs. pSTAT1 and pSTAT4 were assessed by flow cytometry in naive (CCR7^+^CD45RA^+^) or memory (non CCR7^+^CD45RA^+^) CD8^+^ T cells within human PBMCs exposed to IFN-α_2_, β, or ω (at 100 ng/ml) for indicated times. Representative histograms of pSTAT1 and pSTAT4 in CD8^+^ T cell subsets treated with IFN-ω for different times are shown, along with data for all three type-I IFNs on two donor PBMCs. **B**. Representative flow cytometry plots showing tetramer^+^ CD8^+^ T-cells expanded for 10 days upon priming from HLA-A2 donor PBMCs with Melan-A (ELA) peptide (1 µg/ml) alone or in the presence of IFN-α_2_, β, or ω (at 100 ng/ml) (left upper panel) and frequencies of ELA-specific CD8^+^ T cells for 11 donors (lower panel). **C**. Representative flow cytometry plots showing T-bet, Granzyme B and Perforin expression in ELA tetramer^+^ CD8^+^ T cells (upper panel). The dotted line shows the limit for positive expression. Percentage of T-bet, Granzyme B and Perforin positive ELA-specific CD8^+^ T cells (lower panel). Each dot represents one donor (n=11). Horizontal bars indicate median value. Differences between groups were tested using the non-parametric Mann Whitney test.

We then investigated the effect of IFN-ω on the induction of an antigen-specific CD8^+^ T cell response from naive lymphocytes using an *in vitro* model of T cell priming. The Melan-A peptide (ELA) was used as a model antigen owing to the high frequency of specific naive CD8^+^ T cell precursors identified in PBMCs from HLA-A*0201^+^ (HLA-A2^+^) individuals, which can be readily primed *in vitro* from unfractionated PBMCs^29^. This assay enables us to evaluate the effects of IFN-ω on the expansion and functional properties of antigen-specific CD8^+^ T cells 10 days after priming. The expansion of ELA tetramer^+^ CD8^+^ T cells was rather reduced in the presence of IFN-β and IFN-α_2_, but not IFN-ω (**Figure 3B**). IFNs-I, especially IFN-ω, significantly increased the expression of the transcriptional factor T-bet, known to promote CD8^+^ T cell effector functions, such as the secretion of IFN-γ (**Figure 3C**). Moreover, significantly higher levels of the cytotoxins Granzyme B and Perforin were observed in the presence of IFNs-I (**Figure 3C**). Therefore, IFN-ω appears particularly potent at promoting the acquisition of effector functions by CD8^+^ T cells upon priming, without blunting their expansion. These data indicate that IFN-ω can contribute to the induction of strong cytolytic CD8^+^ T cell responses in the context of viral infections.

## Discussion

Altogether, our findings indicate that human IFN-ω is produced by monocytes or pDCs at levels comparable to those of IFN-α_2_ or IFN-β, and can contribute significantly to the first line of defense against viral infections, as well as the induction of potent antiviral CD8^+^ T cell responses. The biological properties of human IFN-ω closely resemble those of IFN-α_2_, including its antiviral activities and immunostimulatory effects. Considering the critical role of IFN-α_2_ or IFN-β in clinical practice, IFN-ω could therefore be considered an important candidate in immunotherapies. Although IFN-ω shares 62% and 33% of its amino acid sequence with IFN-α_2_ and IFN-β respectively, antibodies specific for these two IFNs-I do not cross-react with IFN-ω^30,31^. IFN-ω represents therefore an interesting secondary option for patients who are resistant to IFN-α_2_ or IFN-β due to the formation of autoantibodies during treatment^32^. The lack of *in vivo* mouse models has slowed the development of therapeutic applications using IFN-ω. However, the presence of IFN-ω reduced the production of HbeAg and HBV-DNA synthesis by half in human HBV infected hepatoma cells^33^. Furthermore, IFN-ω appeared to be more potent than IFN-α_2_ in suppressing influenza infection in A549 cell line^34^. Recombinant human IFN-ω was also capable of subduing mRNA levels of HPV11 E6 in the HaCaT cell line^35^. Taken together, these data imply that IFN-ω could exert important antiviral effects on a variety of viruses, not limited to retroviruses as demonstrated in feline models. The study of IFN-ω in clinical trials for the treatment of viral infections should be a priority.

## Materials and methods

(Detailed protocols are provided in supporting information)

### Cells and viruses

PBMCs were obtained from healthy volunteers attending the Etablissement Français du Sang. Jurkat dual cells were obtained from Invivogen and Calu-3 cells from the ATCC. The SARS-CoV-2 strain BetaCoV/France/IDF0372/2020 was supplied by the National Reference Centre for Respiratory Viruses.

### Cytokine quantification and flow cytometry

IFNs-I were detected using specific ELISA kits (Mabtech and eBioscience). For phosphoflow assays, cells were stained with anti-STAT1-AF647 and anti-STAT4-PE (BD Biosciences). *In vitro* priming of antigen-specific CD8^+^ T cells was performed as described previously^36^. Data were acquired using an LSR Fortessa (BD Biosciences) and analyzed with FlowJo (Tree Star Inc.).

### Statistical Analysis

Statistical significance was determined using unpaired T-test with Mann-Whitney (GraphPad Software).

## Acknowledgements

We thank all donors for participating in this study. We grateful to David Price and Sian Sian Llewellyn-Lacey from Cardiff University School of Medicine, Cardiff CF14 4XN, UK for providing peptide-MHC tetramer reagents, and to Atika Zouine and Vincent Pitard for technical assistance at the Flow cytometry facility, CNRS UMS 3427, INSERM US 005, Univ. Bordeaux, F-33000 Bordeaux, France.

## Author Contributions

Conceptualization, V.A., L.P., H.O.N; methodology, V.A., H.O.N, P.P., M-L.A.; investigation and data curation, H.O.N., P.P.; writing—original draft preparation, H.O.N., writing—review and editing, V.A., L.P., P.P., M-L.A; visualization, H.O.N and P.P.; supervision, V.A.; funding acquisition, V.A. All the authors have read and agreed to the published version of the manuscript.

### Abbreviations

IFN: Interferon
IFNs-I: Type-I Interferons
HBV: Hepatitis B virus
HCV: Hepatitis C virus
ISGs: Interferon-stimulated genes
TCR: T cell receptor
pDCs: plasmacytoid dendritic cells
SARS-COV2: Severe acute respiratory syndrome coronavirus 2
STAT1: Signal transducer and activator of transcription 1
STAT4: Signal transducer and activator of transcription 4
PBMCs: Peripheral blood mononuclear cells
PDL1: programmed cell death 1 ligand
rFeIFN: recombinant feline Interfon
rhIFN: recombinant human IFN

## Availability of data and materials

The original contributions presented in the study are included in the article. Further inquiries can be directed to the corresponding author.

